# A novel stabilization mechanism accommodating genome length variation in evolutionarily related viral capsids

**DOI:** 10.1101/2023.11.03.565530

**Authors:** Jennifer M. Podgorski, Joshua Podgorski, Lawrence Abad, Deborah Jacobs-Sera, Krista G. Freeman, Colin Brown, Graham Hatfull, Antoni Luque, Simon J. White

## Abstract

Tailed bacteriophages are one of the most numerous and diverse group of viruses. They store their genome at quasi-crystalline densities in capsids built from multiple copies of proteins adopting the HK97-fold. The high density of the genome exerts an internal pressure, requiring a maturation process that reinforces their capsids. However, it is unclear how capsid stabilization strategies have adapted to accommodate the evolution of larger genomes in this virus group. Here we characterized a novel capsid reinforcement mechanism in two evolutionary-related actinobacteriophages that modifies the length of a stabilization protein to accommodate a larger genome while maintaining the same capsid size. We used cryo-EM to reveal that capsids contained split hexamers of HK97-fold proteins with a stabilization protein in the chasm. The observation of split hexamers in mature capsids was unprecedented, so we rationalized this result mathematically, discovering that icosahedral capsids can be formed by all split or skewed hexamers as long as their T-number is not a multiple of three. Our results suggest that analogous stabilization mechanisms can be present in other icosahedral capsids, and they provide a strategy for engineering capsids accommodating larger DNA cargoes as gene delivery systems.

**Significance Statement:** How capsids are stabilized and change size is an important part of understanding how to design protein containers and understand viral evolution. We describe a novel capsid stability mechanism that allows the capsid to package a larger genome without changing the capsid architecture and have predicted other capsids using this mechanism. Beyond the evolutionary implications, our findings provide a mechanism to increase the amount of DNA packaged in a capsid, offering a solution to engineer gene delivery systems with larger DNA content, a pressing challenge in gene therapy.

## Introduction

Viruses protect their genome in a protein shell called a capsid, which must remain stable in the environment to allow a virus to infect a new host and continue its viral life cycle^1^. One of the most abundant capsids in the virosphere are the ones associated with tailed bacteriophages (or tailed phages). These capsids have adapted to infect bacterial hosts across all environmental conditions^2^ while storing the double-stranded DNA viral genome at the highest density known among biological entities^3^. This condensed viral genome generates 30-50 atmospheres of internal pressure pushing out onto the capsid, and this pressure helps initiate the translocation of the DNA to infect the host^4^. Despite this internal pressure, tailed phage capsids have evolved in size to accommodate from 10 kilo base pairs (kb) to some of the largest genomes found in the virosphere, around 750 kb^5^. A key factor that has allowed tailed phages to achieve this feat is the sophisticated maturation process that stabilizes their capsids. However, there has not been an empirical observation showing the adaptation of a specific stabilization mechanism to accommodate the evolution of a larger genome.

The assembly of tailed phage capsids starts with scaffolding proteins and major capsid proteins (MCPs) nucleating around a protein portal complex, forming an empty shell called the procapsid. The DNA is then packaged through the portal, causing the scaffolding proteins to be either ejected or degraded and the MCPs to undergo conformational changes leading to the final mature polyhedral capsid^6^. This maturation process is possible thanks to the plasticity of the HK97-fold present in the MCPs across tailed phages and other evolutionary related viruses that store their genome at high densities, like tailed archaeal viruses and herpes-like viruses^7 8 9^. The HK97-fold MCPs organize in clusters (capsomers) of five proteins (pentamers) at the vertices of the polyhedral capsid and clusters of six proteins (hexamers) connecting the vertexes. In the maturation process, the MCP hexamers experience the most dramatic conformational changes of any element in the capsid, transitioning often from smaller and skewed hexamers in the procapsid to larger, highly symmetrical hexamers in the mature capsid. This transition introduces stabilization mechanisms like covalent and non-covalent cross-linking of the MCPs^10,11^. It also helps accommodate accessory proteins (also known as cement, reinforcement, and decoration proteins) that can further stabilize the capsid, especially at the local three-fold axes of the icosahedral capsids^12^. It is likely that these stabilization mechanisms have facilitated the evolution of larger viral genomes and capsids. However, the cross-linking and accessory protein stabilization mechanisms characterized in different model systems are too evolutionary distant to reveal the underlying molecular mechanisms able to accommodate an incremental increase in the genome size of the capsid.

To explore this gap, we concentrated on phages infecting the Actinobacteria phylum. These so called actinobacteriophages represent the largest curated collection of viruses with 4,000 annotated phage genomes (as of April 2023)^13^. We recently used a cryo-EM multiplex method to obtain medium-resolution maps (about 6 Å) of multiple capsids^14^. Combining the actinobacteriophages genome nucleotide similarity^15^ and the folded structure of major capsid proteins, we built a structural dendrogram of the actinobacteriophage major capsid proteins^16^ to facilitate navigating evolutionarily related capsids. One of the most remarkable findings from the medium resolution maps was Patience, which was clustered in the Structural Group 11 (or Patience-like). Phage Patience displayed a T=7 extended icosahedral capsid containing 415 HK97-fold MCPs and 420 minor proteins organized in trimers^14^. An intriguing feature of Patience was that its hexamers seem to display a large chasm along their 2-fold local symmetry line, effectively separating the hexamer into two halves. Split or skewed hexamers are common in procapsids but were not previously observed in mature capsids, which usually contain 6-fold symmetry hexamers. Additionally, the chasm had an apparent density within it that could not be resolved. This density represented a candidate for a novel reinforcement mechanism that could change the established paradigm of mature capsids. Thus, in this study, we followed up that preliminary observation to obtain a high-resolution cryo-EM map of Patience to 2.4 Å as well as the map for an evolutionary-related phage, Adjutor, which displayed structural protein homologies while storing a different genome length. Our expectation was that we could confirm the observation of skewed mature hexamers with a new reinforcement mechanism and capture how the same reinforcement mechanism adapted to accommodate a different genome length.

The reconstruction of the high-resolution cryo-EM maps for Patience and Adjutor confirmed the existence of split hexamers containing an accessory protein that stabilizes the hexamers and has adapted its length to accommodate the change in genome length without having to increase the size of the capsid. The derivation of a new mathematical theorem allowed us to predict what other capsid architectures could use an analogous stabilization mechanism that could adapt to increments of genome length.

## Results and Discussion

### Novel accessory protein interacting with skewed mature hexamers

The high-resolution cryo-EM map of Patience to 2.4 Å confirmed the presence of split hexamers (Fig. 1A, Table S1 for final resolution and collection parameters) and revealed that the chasm’s density was associated with a different protein from the major capsid protein (Fig. 2). *De novo* model building of the density identified the protein as gp4. Our annotation was further supported by previous mass spectrometry analysis of the Patience capsid, which had identified gp4^17^ within the mature viral capsid. Gp4 (Fig. S1) is a small 102 amino acid long protein of 10.7 kDa that forms a dimer within the hexamer and is only observed in the hexamer (the pentamer did not show any evidence of a density that could be gp4). We note that there was some density we could not model under the local 3-fold axes (Fig. S2) that showed a small helix but at too low a resolution to model. Three gp4 dimers organized around the global 3-fold axis forming a three-point star complex (trimer of dimers) that does not interact with other three point star gp4 complexes, nearby pentamer major capsid proteins, or the nearby minor capsid proteins, which sits on top of the major capsid proteins at the local 3-fold axes (Fig. 1A). Thus, gp4 only interacts with the major capsid proteins of the hexamer. Circular dichroism analysis (Fig. S3) of unbound Patience gp4 showed that the secondary structure of the protein is a random coil, while in the capsid it forms a helix.

**Figure 1.**
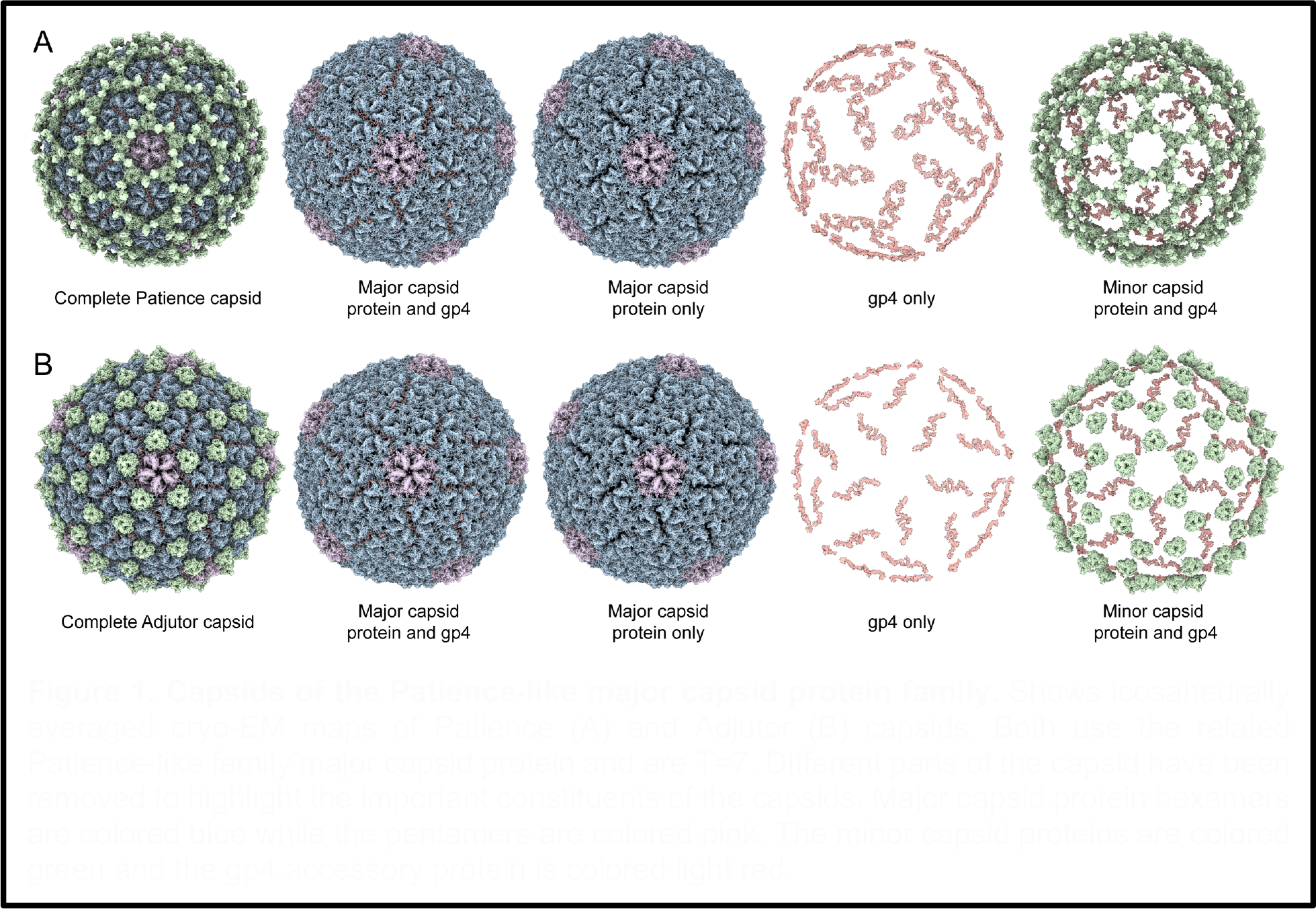
Capsids of the Patience-like major capsid protein family. Shows icosahedrally averaged cryo-EM maps of Patience (A) and Adjutor (B) capsids. Both use the related Patience-like family major capsid protein and are T=7. Different parts of the capsid have been removed to highlight the important constituents of the capsids. Major capsid protein hexamers are colored blue while the pentamers are colored pink. The minor capsid proteins are colored green and the gp4 accessory protein is colored light red.

**Figure 2.**
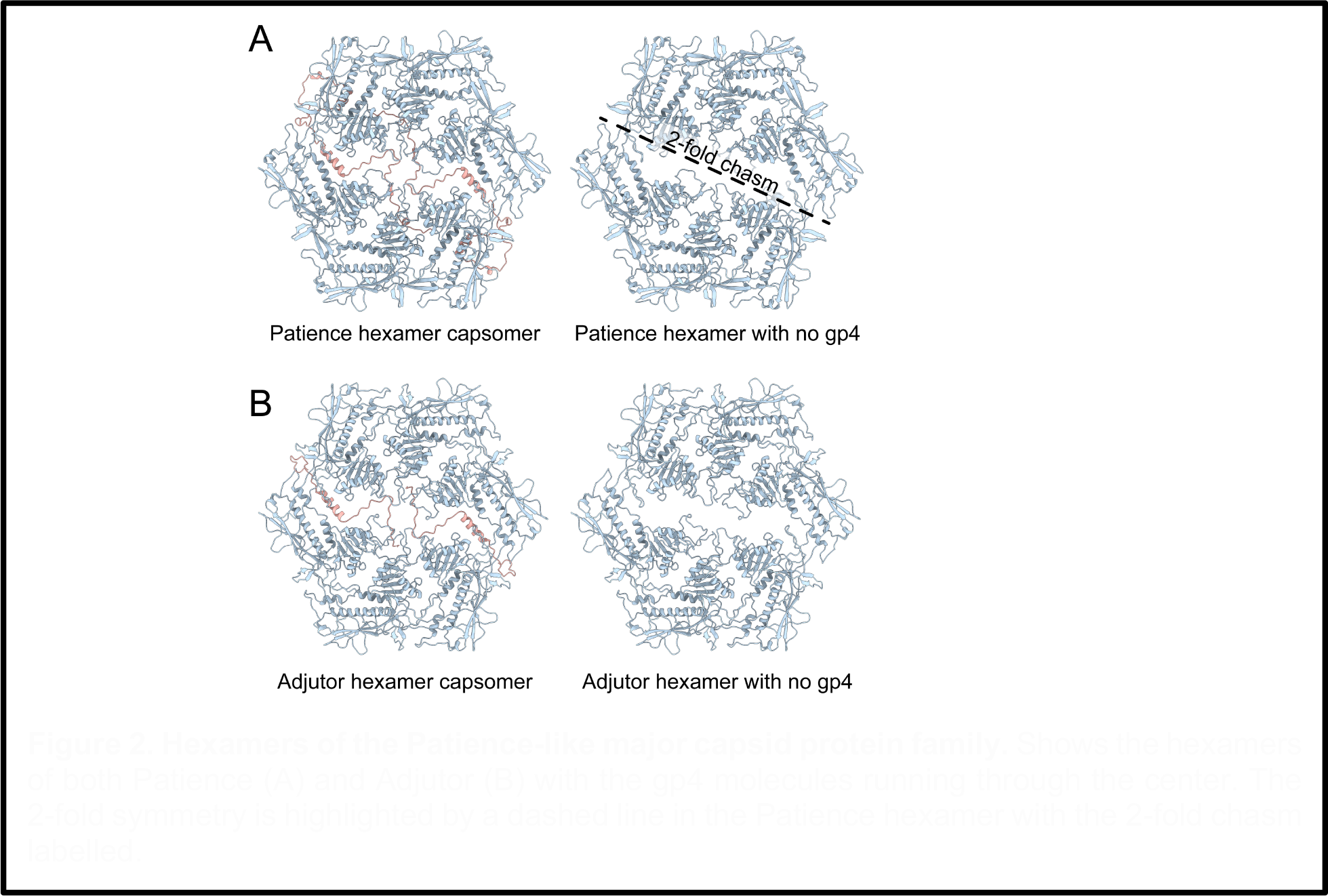
Hexamers of the Patience-like major capsid protein family. Shows the hexamers of both Patience (A) and Adjutor (B) with the gp4 molecules running through the center. The 2-fold symmetry is highlighted by a dashed line in the Patience hexamer with the 2-fold chasm labelled.

### Patience-like phages show a conserved stabilization mechanism

The search of gp4 homologs among actinobacteriophages yielded candidates in the AC, D, H and R clusters (Table S2). The predicted protein structure with Alphafold revealed that the candidate proteins in the D and H clusters adopted a similar structure to Patience gp4. We decided to focus on Adjutor (D cluster, gp4 homolog to Patience gp4) to carry out cryo-EM to confirm the Alphafold prediction (Fig. 1B) because it had the smallest gp4 homolog.

The cryo-EM map showed that Adjutor has a similar capsid morphology to Patience. The minor capsid protein lacks the peanut-like protein domain found in Patience. However, the rest of the minor capsid protein of Adjutor is structurally very similar to Patience’s (Fig. S1) and is part of the same protein family. The cryo-EM map was at a high enough resolution (2.7 Å, Table S1 for collection parameters) to once again identify the protein density found between the two halves of the major capsid protein hexamer as gp4 of Adjutor. Gp4 of Adjutor is half the size of Patience gp4 at just 54 amino acids and a molecular weight of 6.2 kDa. Once again, it exists as “dimers” within the hexamer and “trimers” around the global 3-fold axes. (Fig. 1B). As observed in Patience, Adjutor-gp4 only contacts the major capsid protein hexamer capsomer and makes no contact with the pentamer or minor capsid protein. Likewise, no density for Adjutor-gp4 was observed in the pentamer capsomer.

### Gp4 bridging of split hexamers allows an increased size of genome to be packaged

The smaller size of the Adjutor-gp4 proteins suggests that it is a simpler version of the Patience-gp4. A single Adjutor-gp4 protein (Fig. 3B) makes hydrogen bonds and salt bridges (Table S3, Fig. 3B) to four of the chains of the hexamer (A, B, E, and F). Most of the buried surface area in a single Adjutor-gp4 is between chains A and F, i.e., the two major capsid proteins on either side of the 2-fold chasm. The second Adjutor-gp4 protein (Fig. 3B) performs the same function but for the opposite end of the hexamer 2-fold chasm, whereby it makes the most contacts with chains C and D. The two adjutor-gp4 chains fill all the space of the chasm, with the weaker contacts presumably stabilizing the position of the gp4 molecules.

**Figure 3.**
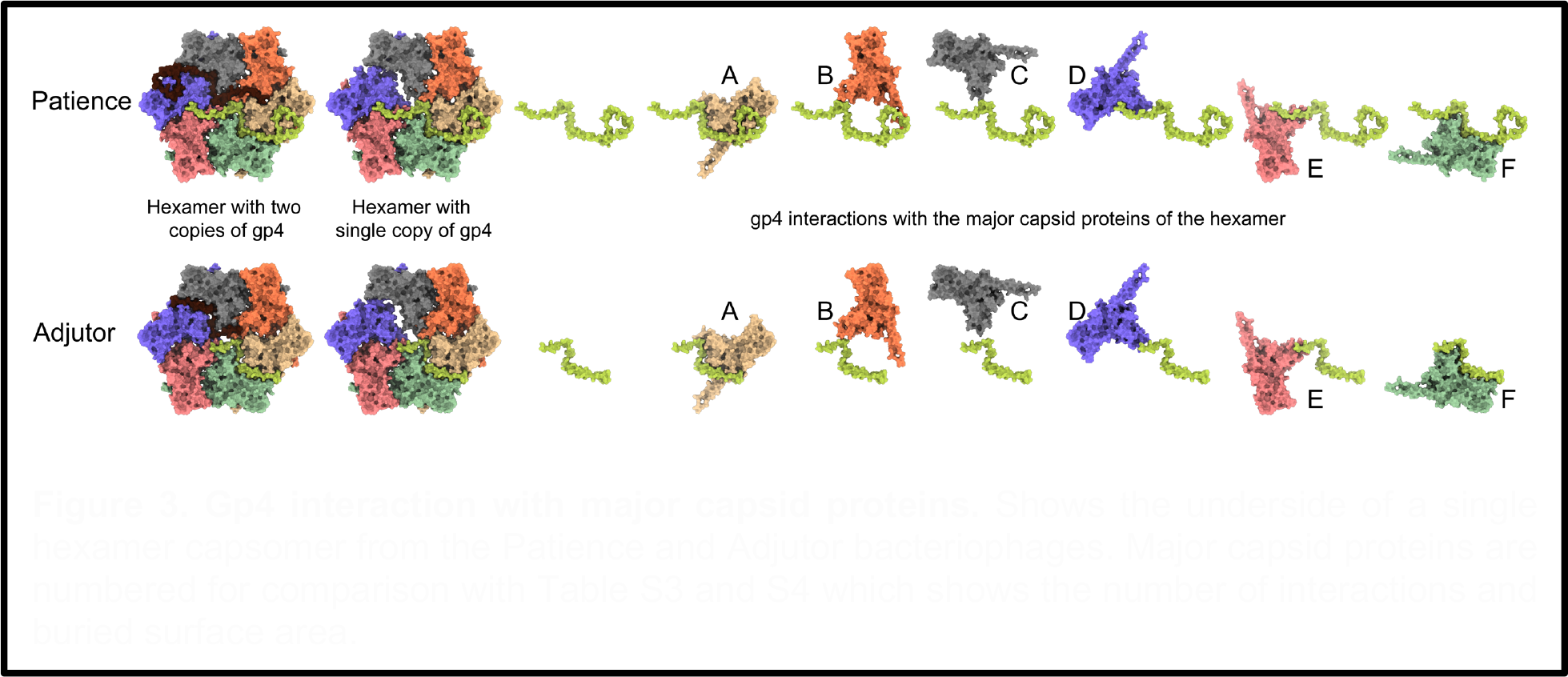
Gp4 interaction with major capsid proteins. Shows the underside of a single hexamer capsomer from the Patience and Adjutor bacteriophages. Major capsid proteins are numbered for comparison with Table S3 and S4 which shows the number of interactions and buried surface area.

The Patience-gp4 makes far more contacts than the Adjutor-gp4, and a single Patience-gp4 (Fig. 3A) makes interactions (both hydrogen bonds/salt bridges and buried surface area) with every copy of the major capsid protein in the hexamer (Table S4). Like with Adjutor-gp4, most interactions are with the A and F chains of the hexamer (with the second Patience-gp4 molecule having the most interactions with chains C and D). In contrast to Adjutor-gp4, Patience-gp4 is far longer and threads its way between the spaces of the major capsid proteins at its N and C termini. At the C-terminal end, this results in a peg-in-hole interaction, with the thirty-five residues between Patience-gp4 60 and 93 forming a loop structure (the hole) around residues 335-345 of chain A major capsid protein, with a small helix turn (339-341) acting as the peg (Fig. S5). Several salt bridges and hydrogen bonds stabilize the interaction. At the N-terminal end of Patience-gp4, it continues to thread between Chain E and Chain F of the major capsid proteins, making several stabilizing interactions with those proteins. The major capsid proteins of Adjutor have similar pockets that could accommodate a longer gp4, and Adjutor-gp4 follows the same path as Patience-gp4 but lacks the length to thread between the other major capsid proteins.

Patience and Adjutor display the same internal capsid diameter (63 nm), but Patience packs a larger genome (70.5 kb) than Adjutor (64.5 kb). This constitutes a 10% increase in genome density, which in tailed phages implies an increase of internal pressure^18,19^. The concomitant increase of the gp4 length and number of interactions in Patience with respect to Adjutor, was attributed to this increase in internal pressure. We note that the genome of Adjutor is approximately 30% larger than other well studied T=7 phages, for example lambda, HK97 and L5. This suggests that gradual changes in gp4 length would help accommodate slight genome length increases in the evolution (or engineering) of tailed phages. This mechanism, however, requires the formation of split hexamers, which had not been observed in mature capsids previously. To show that all split hexamers (and, therefore, analogous stability mechanisms) are potentially relevant to other capsids, we investigated mathematically the different icosahedral capsid architectures that can contain all-split hexamers. This mathematical prediction is shared in the last section of the results. Before presenting those results, it is worth completing the experimental characterization of the structural novelties of Patience and Adjutor.

### The E-loop is over-extended in the major capsid proteins that bridge the 2-fold chasm

The two chains in Patience and Adjutor (A and D) have E-loops that bridge the 2-fold chasm (Fig. 3). This is achieved by straightening and elongating the E-loop starting at Val73 (Fig. S6). In the other four chains (B, C, E, and F), the E-loop folds into a U-shape before forming the two beta strands starting at Val58. In the A and D chains, the two beta strands are abolished, and the U-shape is straightened out, leading to the elongation of the E-loop. The elongated E-loops are stabilized by an intramolecular salt bridge between Glu78 and Arg370, with the arginine and glutamic acid repositioning to make the interaction. In the other four chains, these amino acids are pointed away from one another, and Glu78 is part of the two beta strands of the E-loop.

The straightening is also supported by the gp4 molecule with five hydrogen bonds between gp4 and the E-loop of chains A and D. In Patience, the E-loop is further constrained by the C-terminus of gp4 that folds over the top of the E-loop at the straightened position. This straightening of the E-loop allows the loop region to make the same contacts in A and D with the adjacent major capsid proteins as B, C, E, and F do, despite the increased distance the E-loop must cover. Thus, the first salt bridge between Glu85 (E-loop) and Arg148 of the spine helix and the final salt bridge between Arg98 (E-loop) and Glu26 of the N-arm are spatially equivalent in all chains, with the N-arm folding over the E-loop and locking it into position above the spine helix of an adjacent chain. The same overall mechanism for E-loop extension is observed for Adjutor. The amino acids making hydrogen bonds are different, and Adjutor shows more hydrogen bond/salt bridge interactions than Patience (37 intermolecular bonds of Adjutor E-loop and 21 in Patience E-loop). Despite the decrease in length of gp4 in Adjutor, the C-terminus still folds over the straightened E-loop at the same position as in Patience to lock in the structure.

### The major capsid protein of the Patience-like actinobacteriophages shows variation from the canonical HK97-fold and forms a putative metalloprotein

The major capsid protein of Patience and Adjutor shows major differences to the canonical HK97 phage HK97-fold^7^. It has a seven-strand beta-sheet in the A-domain with an alpha helix (211-225) that lies across it. The major capsid proteins display an extended G-loop, forming a small beta-hairpin. Following the G-loop is a loop that we have previously described in other actinobacteriophages (and others have described in phage T7^20^) that make no inter or intra-molecular interactions aside from a possible hydrogen bond between Arg188 and Thr20 of two different chains (but only in chains C and F where the N-arm is pointed up). Unlike most previously characterized HK97-folds in tailed phages, the N-arm does not make long-range interactions with other major capsid proteins. Instead, the N-arm forms a helix that runs almost parallel to the P-domain spine helix below it (Fig. S7) before turning back on itself. At the turn, it makes a single salt bridge with the G-loop of a neighboring major capsid protein (Asp 40 of the N-arm and Gln171 of the G-loop). This interaction only occurs between major capsid proteins located within each half. The interaction is lost for the chains that bridge the 2-fold chasm (Chains C and D or E and F). In these chains (C, D, E, F), Gln171 of the G-loop makes no interaction, while Asp40 interacts with Asn100 of the gp4 chain. After the N-arm turns back on itself, it makes several intramolecular contacts with the P domain before crossing over the E-loop of an adjacent major capsid protein. It ends in between the spine helix and the N-arm helix, pointing down between those two helices (Fig. S7). In chains C and F, the two N-terminal amino acids sit up compared to the other chains; the N-termini of these chains appear to be displaced by the presence of the gp4 molecules that lie adjacent to the N-arm helix.

His198, His200, Glu223, and His224 in the A domain form a putative metal binding domain. There is clear density in the cryo-EM derived map for a metal ion (Fig. S8). We have not modeled a metal ion into that position since it is not practical to definitively identify the metal ion. Bioinformatic analysis using the Metal-Ion Binding^21^ site online server (Table S5) suggested Cobalt and Zinc for Patience and Adjutor, respectively (Fig. S8), although Zinc scored highly for Patience and the related Candle bacteriophage. Other viral proteins that are evolutionarily distinct to the HK97-fold, for example, Hepatitis B virus^22^ is known to use Zinc in its capsid proteins, and this is the most common metal that interacts with viral proteins^23^. We therefore speculate that Zinc is the most likely metal ion to be present in all the Patience-like bacteriophages. The position of the metal binding domain at the bottom of the helix that crosses the beta-sheet in the A-domain suggests that it is involved in locking the bottom of the helix into the correct position. This is likely to be especially important in chains A and D where there 2-fold chasm means that there is no neighboring major capsid protein A domain to stabilize the A-domain helix. The metal binding domain is conserved across the Patience-like actinobacteriophages. We picked a representative major capsid protein from each cluster that uses the Patience-like major capsid protein and predicted their structure using Alphafold. All the major capsid proteins were structurally near identical despite relatively low amino acid sequence identity (Fig. S9). All have a putative metal binding domain at the same position in the major capsid protein and use three histidines but show varying amino acids that are also part of the putative metal binding site (Fig. S10). For example, Adjutor has an additional histidine and glutamic acid, while Candle (cluster R) has five histidines.

### Icosahedral capsids that can be built with split or skew hexamers

To determine if a 2-fold-like chasm mechanism could be relevant in other capsids, we investigated mathematically what icosahedral capsids can contain all-split hexamers. A key property of icosahedral capsids is that they display three types of axes of symmetry: 5-fold, 3-fold, and 2-fold. MCPs sitting on the 5-fold axes form 5-fold symmetric pentamers that generate the topological curvature necessary to close the quasi-spherical capsid^24,25^. However, the 3-fold and 2-fold axes of the capsid do not necessarily pass through the center of a hexamer because the distribution of hexamers, as shown in Fig. 4, changes depending on the icosahedral T-number, which is defined by equation 1.

**Figure 4.**
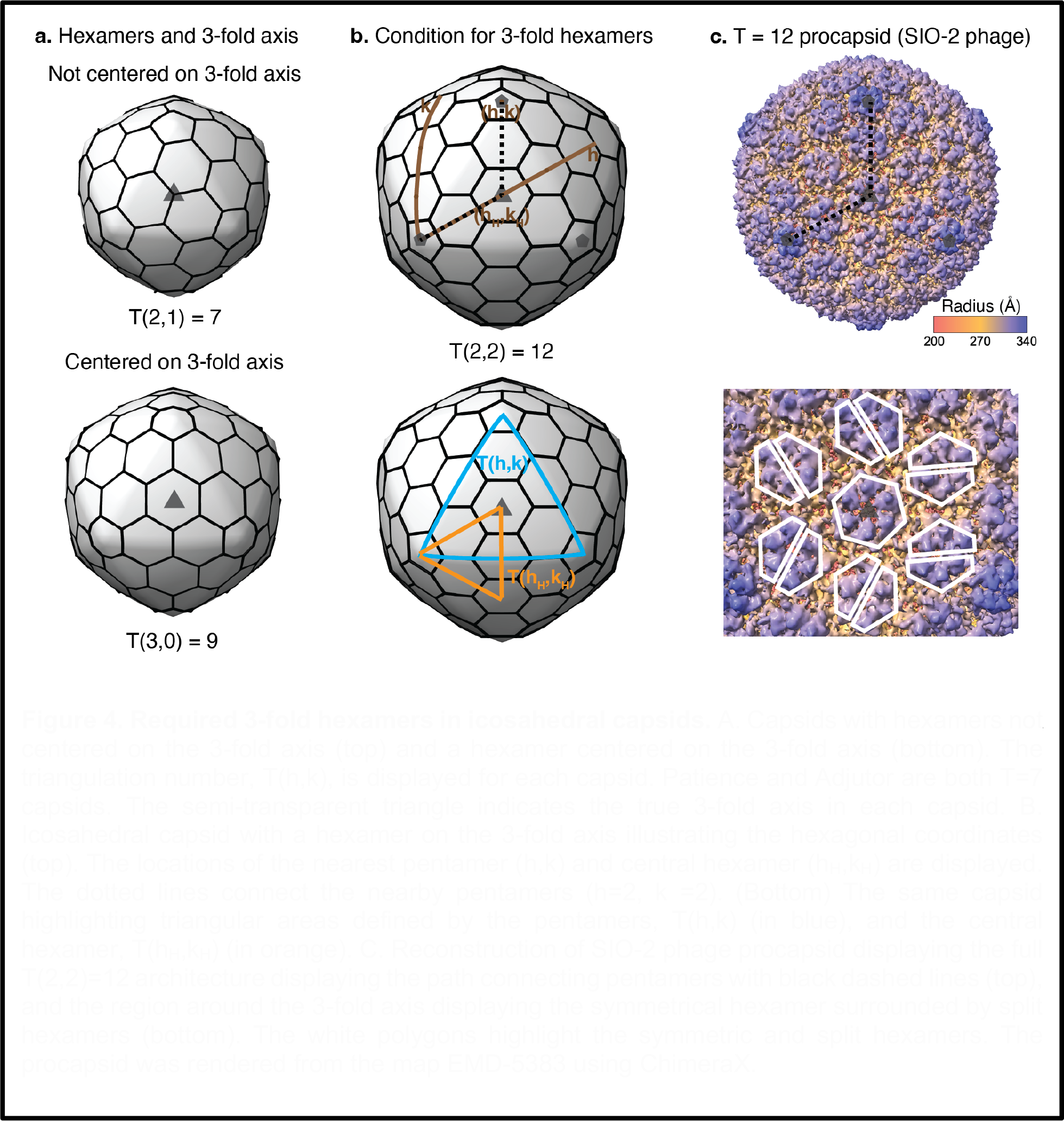
Required 3-fold hexamers in icosahedral capsids. A. Capsids with hexamers not centered on the 3-fold axis (top) and a hexamer centered on the 3-fold axis (bottom). The triangulation number, T(h,k), is displayed for each capsid. Patience and Adjutor are both T=7 capsids. The semi-transparent triangle indicates the true 3-fold axis in each capsid. B. Icosahedral capsid with a hexamer on the 3-fold axis illustrating the hexagonal coordinates (top). The locations of the nearest pentamer (h,k) and central hexamer (h_H_,k_H_) are displayed. The dotted lines connect the nearby pentamers (h=2, k =2). (Bottom) The same capsid highlighting triangular areas defined by the pentamers, T(h,k) (in blue), and the central hexamer, T(h_H_,k_H_) (in orange). C. Reconstruction of SIO-2 phage procapsid displaying the full T(2,2)=12 architecture displaying the path connecting pentamers with black dashed lines (top), and the region around the 3-fold axis displaying the symmetrical hexamer surrounded by split hexamers (bottom). The white polygons highlight the symmetric and split hexamers. The procapsid was rendered from the map EMD-5383 using ChimeraX.

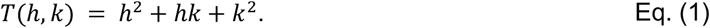

Here, *h* and *k* are the steps necessary on the hexagonal lattice to join two nearby pentamers, and the icosahedral capsid is formed by 60T MCPs distributed in 10(T-1) hexamers and 12 pentamers. Hexamers with 6-fold symmetry guarantee the formation of any icosahedral capsid because they are compatible with both 3-fold and 2-fold symmetry axes, which might explain why this configuration is so common in mature capsids. However, the split hexamers observed in Adjutor and Patience (and the common skew hexamers of tailed phages’ procapsids, Fig. S11) only have 2-fold symmetry. Since this configuration would cause a symmetry mismatch in capsids that use T-numbers where the center of a hexamer sits on a global 3-fold axis, this reduces the problem of finding capsids with all-split (or skew) hexamers to determine mathematically the T-numbers that do not contain a hexamer on the center of a global 3-fold axis.

We initially investigated the range of icosahedral capsids associated with the HK97-fold, that is, from T=1 to T=52, computationally. The generated icosahedral capsids showed eight capsids containing a hexamer sitting on a global 3-fold axis: T = 3, 9, 12, 21, 27, 36, 39, and 48 (Table S6) as illustrated by T=9 and T=12 in Figs. 4A and 4B. The remaining twelve capsids could contain all split/skew hexamers: T= 4, 7, 13, 16, 19, 25, 28, 31, 37, 43, 49, 52, as illustrated by the T=7 capsid in Fig. 4A. Thus, the capsids displaying hexamers on the 3-fold axes shared a mathematical pattern: Their T-numbers were multiple of 3, e.g., T = 21 = 3 × 7 (Table S6).

Interestingly, the remaining factor in those T-number, that is, T/3, recovered the sequence of classic T-numbers, that is, 1, 3, 4, 7, 9, 12, 13, and 16. These observations from the computational analysis let us formulate the following general theorem: “An icosahedral capsid has a hexamer centered on a true 3-fold if and only if the T-number is multiple of three.”

We proved the theorem as follows. In capsids with hexamers centered on a true 3-fold, one can define *h*_*H*_ and *k*_*H*_ as the hexagonal lattice steps joining a pentamer with the hexamer sitting on a global 3-fold axis, as illustrated for T=12 in Fig. 4B. By symmetry, the triangle formed by three pentamers is three times larger than the triangle formed by one pentamer and the two vertices placed on nearby 3-fold axes. The normalized area of the small triangle formed by the central hexamer follows the same expression as the classical T-number sequence given by Eq. (1), except that the steps used are *h*_*H*_ and *k*_*H*_ instead of *h* and *k*. The final triangulation number is three times this small triangle, leading to the T-numbers, *T*_*H*_, for capsids with hexamers centered on the true 3-fold axes:

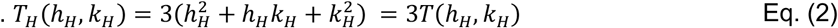

The associated steps h and k can be expressed as a function of *h*_*H*_ and *k*_*H*_ by *h = h*_*H*_ *+ k*_*H*_ and *k = k*_*H*_ *-h*_*H*_ (Fig. 4B). Introducing these expressions in Eq. (1) offers an alternative way to derive *T*_*H*_:

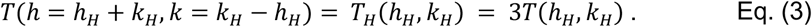

Eqs. (2) and (3) imply that any T-number multiple of three is associated with a capsid with centered hexamers on the global 3-fold axes. This is equivalent to say that any T-number multiplied by three leads to another T-number with hexamers centered on the 3-fold axes. For example, T=1 leads to T=3, T=3 to T=9, T=4 to T=12, etc. This theorem predicts that capsids and procapsids (immature capsids) of tailed phages forming, e.g., T=3, 9, or 12, will display at least 20 hexamers with 3-fold (or 6-fold) symmetry on the global 3-fold icosahedral axes. This prediction was supported empirically by the published reconstruction of the procapsid of SIO-2 tailed phage, which adopts a T=12 capsid with non-split hexamers at the true 3-fold axes^26^ (Fig. 4C).

The theorem derived above can be also stated as follows: “An icosahedral capsid can be formed with all-skew (split or 2-fold) hexamers if and only if the T-number is not multiple of three.” This captures all the capsids that could display an analogous stability mechanism as Patience and Adjutor. Notice that since the T-numbers for centered hexamers are generated as three times a classic T-number, these capsids are less frequent than those with all skey/split hexamers. In particular, on average one expects that 1 of every 3 capsids will require a centered hexamer, while 2 of every 3 can accommodate all skew hexamers. This is consistent with the observation made computationally where about 2/3 of the capsids explored (12 out of the 20) could accommodate all skew/split hexamers while only about 1/3 (8 out of 20) displayed a 3-fold centered hexamer.

In conclusion, we have structurally characterized two actinobacteriophages that use a previously uncharacterized capsid organization which involves split hexamers that are stabilized by a novel accessory protein. Bacteriophages are ancient and have had a long time to explore the multiple solutions to create an icosahedral capsid; these two phages reveal one such solution. The capsid organization allows the phage to package larger genomes without having to change the T-number. We predict that capsid architectures that can accommodate all skew/split hexamers and benefit from the chasm-stabilization mechanism described here are characterized by T-numbers that are not a multiple of three due to symmetry constraints.

## Materials and Methods

### Production and purification of Bacteriophages for Cryo-Electron Microscopy

Phages were made as previously described^16^. Each bacteriophage was mixed with their respective host (Table S7), plated in top agar on Luria agar plates, and incubated overnight at the temperatures shown in Supporting Information Table 1 to produce twenty webbed plates. Webbed plates were flooded with 5 mL of Phage Buffer (10 mM Tris-HCl pH 7.5, 10 mM MgSO_4_, 68 mM NaCl, 1 mM CaCl_2_) and incubated overnight at room temperature to allow diffusion of the phages into the Phage Buffer. This solution was aspirated from the plates and centrifuged at 15,000× *g* for 15 min at 4 °C to remove cell debris. Phage particles were initially pelleted using an SW41Ti swinging bucket rotor (Beckman Coulter, Brea, CA) at 30,000 rpm for 3 hours using 12.5 mL open-top polyclear tubes (Seton Scientific, Petaluma, CA). The phages in the pellet were resuspended in 4 mL of Phage Buffer by rocking overnight at 4 °C. To further purify the phage particles in this solution, isopycnic centrifugation was performed by adding 5.25 g of CsCl to the 7 mL of phage lysate. The CsCl/phage solutions were centrifuged at 40,000 rpm in an S50-ST swinging bucket rotor (Thermofisher Scientific, Waltham, MA) for 16 h, and the phage particle band (that appeared roughly halfway down the tube) was removed with a syringe and needle (around 1 mL removed per band). The phage particles were then dialyzed three times against Phage Buffer to remove CsCl. This was accomplished by placing the CsCl/phage solution in dialysis tubing with a 50 kDa MWCO. The phages were pelleted at 75000 rpm in an S120-AT2 fixed angle rotor (Beckman Coulter, Brea, CA) to allow for concentration. The phage particles were resuspended in 20 μL of Phage Buffer with gentle pipetting.

### Preparation of Cryo-Electron Microscopy Grids

Three microliters of concentrated phage particles (around 10 mg/mL) were added to plasma-cleaned Au-flat 2/2 (2 μm hole, 2 μm space) cryo-electron microscopy grids (Protochips, Morrisville, NC, USA) using a Vitrobot mk IV (FEI, Hillsboro, OR, USA). Grids were blotted for 5 s with a force of 5 (a setting on the Vitrobot) before being plunged into liquid ethane.

### Cryo-Electron Microscopy

Data were collected on a 300 keV Titan Krios (FEI, Hillsboro, OR, USA) at the Pacific Northwest Center for Cryo-EM with a K3 direct electron detector (Gatan, Pleasanton, CA, USA). Table S1 provides the collection parameters for each phage.

### Cryo-Electron Microscopy Data Analysis

Relion 3.1.1^27^ was used for phage capsid map reconstructions using the standard workflow. CTF Refinement was performed using the default settings. Ewald sphere correction was performed for each particle using the relion_image_handler command included with Relion. The mask_diameter value used in the Ewald sphere correction is reported in Table S1.

### Model building

The amino acid sequence of the major capsid protein was folded using Alphafold version 2.0^28^ using the default settings on a local device. The highest-ranked prediction model was evaluated and fitted into the cryo-EM map using the “Fit in Map” command in ChimeraX version 1.3^29^. To further fit the density, Coot version 0.9.2^30^ was used to fit the model using the “Stepped sphere refine active chain” provided by the Python script developed by Oliver Clarke^31^. Following this, any outlying parts of the protein chain were manually moved to density. The real-space refinement tool of Phenix version 1.19.2-4158^32^ was used with default settings to refine the model, and Coot was used to manually fix issues identified in the Phenix refinements. This was done iteratively until the final step of using the ChimeraX plugin, Isolde version 1.3^33^, to refine the major capsid protein model. The whole model simulation temperature was set to 20 degrees Kelvin, and default parameters were used. After the first major capsid unit was finished, the process was repeated to make up the asymmetric unit, with a final Isolde refinement.

### Purification and circular dichroism of recombinant gp4

The Patience gp4 construct was designed with a N-terminal MBP tag separated from the sequence with a factor Xa protease cleavage site and a C-terminal Hexa-His tag separated from the sequence by a TEV protease cleavage site. The DNA fragment encoding the protein was codon optimized for *E. coli* protein expression and synthesized by GenScript, USA before being cloned into the pETDuet-1 vector. The protein was produced in NEBExpress Competent *E. coli* High Efficiency cells (New England BioLabs, USA). The cells were grown in luria broth media containing 3% glucose and 100 μg/mL or ampicillin at 30°C until the O.D. reached 0.6 then induced with 1 mM IPTG and grown for 3 hours at 30°C. The cells were pelleted by centrifugation at 15,000xg for 15 minute and frozen at −80°C. The frozen cell pellets were resuspended in lysis buffer (50 mM TRIS pH 7, 150 mM NaCl, 1 mM EDTA, Roche’s complete Protease inhibitors, 1% Triton X-100, 1 mM BME, and small quantities of DNase and RNase). The resuspended cells were ruptured by the One-Shot Cell Disruptor (Constant Systems, UK) and the lysate was incubated for 1 hour shaking at room temperature. The lysate was centrifuged at 30,000xg for 30 minutes at 4°C to remove cell debris. The lysate was run over a MBP column (MBPTrap HP Cytiva, USA), dialyzed into Ni buffer A, run over a Ni-NTA affinity column (HisTrap Cytiva, USA), and dialyzed back into MBP buffer A. MBP buffer A is 50 mM TRIS pH 7, 150 mM NaCl, 1 mM EDTA, and 0.1% Triton X-100 and MBP buffer B has 10 mM maltose added. Ni buffer A is 50 mM TRIS pH 7, 150 mM NaCl, and 0.1% Triton X-100 and Ni buffer B has 200 mM imidazole added. The fraction containing protein was then cut for 48 hours at 4°C with 1 mg of TEV protease per 100 mg gp4 and 1 mg of factor Xa protease per 100 mg gp4. The cut fraction was run on a Superdex 75 Increase 10/300 GL using 1/3x PBS buffer (Cytiva, USA). The protein-containing fraction was concentrated in a 3 kDa MWCO Amicon until 3 mg/mL. The CD spectra was measured from 180 nm to 280 nm at 4°C.

### Data Deposition

Protein databank accession numbers are as follows:

Patience: PDB 8GIU, EMD-40077, EMPIAR-11498.

Adjutor: PDB 8SAJ, EMD-40271, EMPIAR-11499.

### Computationally generated icosahedral capsids

Icosahedral architectures within the range of structures observed in shells formed by HK97-fold capsid proteins^34^, that is, T=1 to T=52, where the T-number or triangulation number follows the Diophantine equation^24,25^ shown in Eq. (1) and described in the text right after.

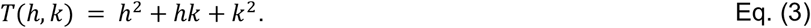

The equation was implemented on a python script. The T-numbers were generated using *h* and *k* values, satisfying *T(h,k) ≤ 52*. These capsids were rendered in 3D using the hkcage tool from ChimeraX^35,36,29^. Images centered on a 3-fold axis were saved for analysis (see Table S6). The process was automated developing a ChimeraX script in cxc file format. The instructions and codes to generate the table and 3D renders for the analysis are publicly available in the Zenodo repository under the digital object identifier (DOI): https://doi.org/10.5281/zenodo.8237508.

## Acknowledgments

We thank the faculty and students in the PHIRE, SEA-PHAGES and the Mycobacterial Genetics Course (2009-2010) programs for their efforts in phage isolation and characterization.

Funding: A.L. National Science Foundation (Award #1951678); NIH grant GM131729 and HHMI grant GT2053 to GFH.

