## Supporting figures and tables for "A novel stabilization mechanism accommodating genome length variation in evolutionarily related viral capsids"

### Supporting Information for “A novel stabilization mechanism accommodating genome length variation in evolutionarily related viral capsids”

**Authors:** Jennifer M. Podgorski[1], Joshua Podgorski[1], Lawrence Abad[2], Deborah Jacobs-Sera[2], Krista G. Freeman[2], Colin Brown[3,4], Graham Hatfull[2], Antoni Luque[3,5,6,7]* Simon J. White[1]*

**Affiliations**

**1.** Biology/Physics Building, Department of Molecular and Cell Biology, University of Connecticut, 91 North Eagleville Road, Unit-3125. Storrs, CT 06269-3125, USA

**2.** Department of Biological Sciences, University of Pittsburgh, 4249 Fifth Avenue, Pittsburgh, PA 15260, USA

**3.** Viral Information Institute, San Diego State University, 5500 Campanile Drive, San Diego, CA 92182, USA

**4.** Department of Physics, San Diego State University, 5500 Campanile Drive, San Diego, CA 92182, USA

**5.** Department of Mathematics and Statistics, San Diego State University, 5500 Campanile Drive, San Diego, CA 92182, USA

**6.** Computational Science Research Center, San Diego State University, 5500 Campanile Drive, San Diego, CA 92182, USA

**7.** Department of Biology, University of Miami, 1301 Memorial Dr, Coral Gables, FL 33155, USA

* Authors to whom correspondence should be addressed.

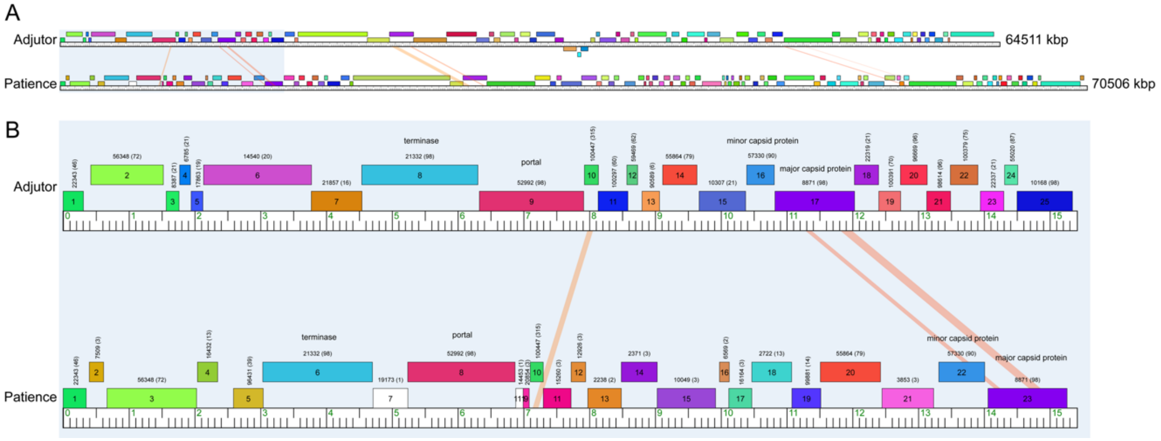


**Figure S1. Genomes of Patience and Adjutor.** A) shows the full genome of Adjutor and Patience. Orange/yellow lines between the two genomes shows genes that have nucleotide sequence identity. The length of the genomes is shown to the right. B) shows the zoomed in area of the structural genes. Certain genes are annotated. The number above each gene is the pham number. The number in brackets is the number of other phages with the same pham protein. For example, the major capsid protein is Pham 8871 with 98 other bacteriophages containing the same Pham protein. Genome cartoon was created with Phamerator[1].

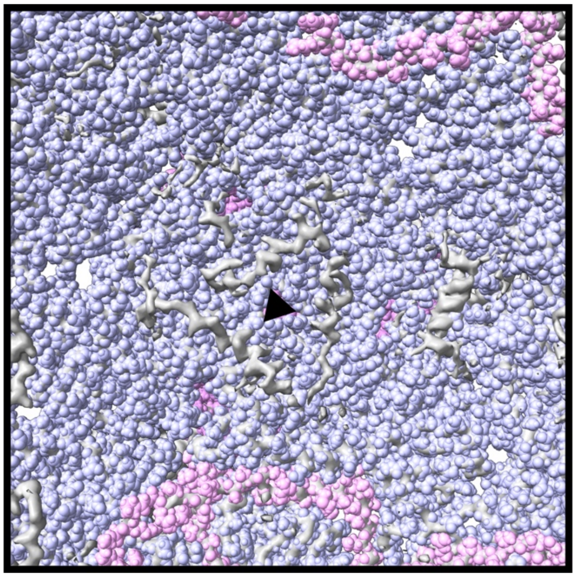

**Figure S2. Unmodelled density in Patience.** The areas in grey are unmodelled density observed on the underside of the capsid near the local three-fold (black triangle) axis. Blue shows major capsid protein while pink shows gp4. The helix can be observed on the right hand side.

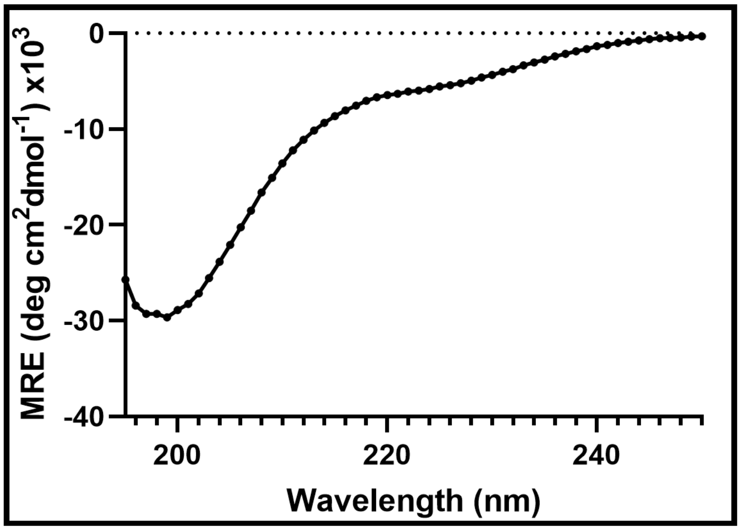


**Figure S3. Free Patience gp4 exists as a random coil.** A CD spectra of free Patience gp4 at 1 mg/mL.

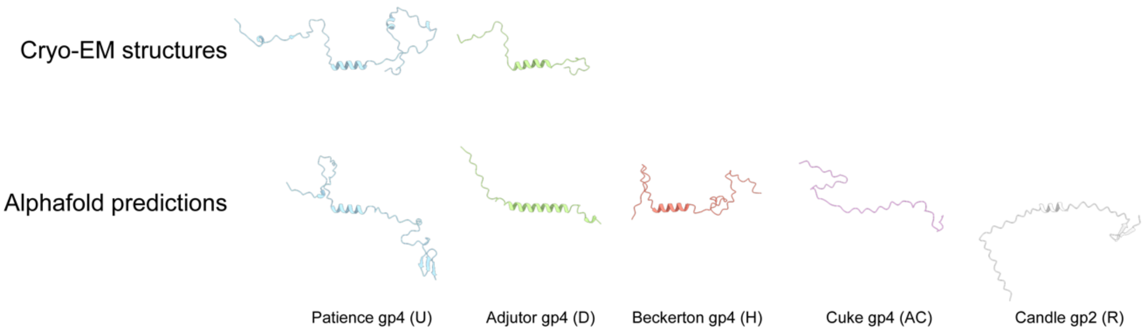


**Figure S4. Alphafold predicted structures of putative gp4 homologs.** Patience and Adjutor cryo-EM models and their predictions are shown for comparison.

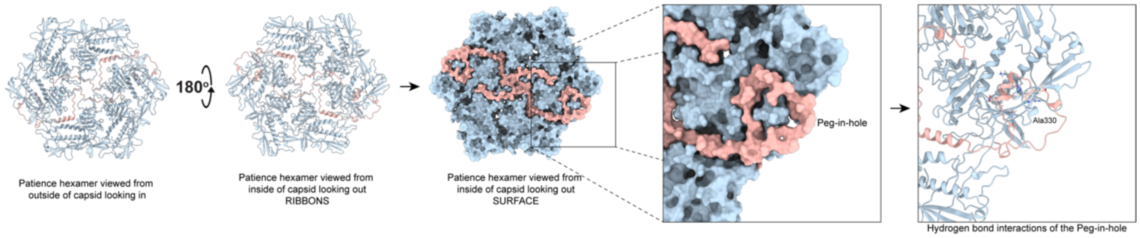


**Figure S5. Peg-in-hole interaction of Patience gp4.** Shows the ribbon diagram of the Patience hexamer capsomer from the outside looking into the capsid. This is in flipped 180o so that the underside of the ribbon diagram of the hexamer is seen. The surface is then shown and zoomed in to highlight the peg-in-hole interaction. Finally, the ribbon diagram of the peg-in-hole is shown with Ala330 labelled as the center of the major capsid protein peg. The hydrogen bonds and salt bridges are revealed.

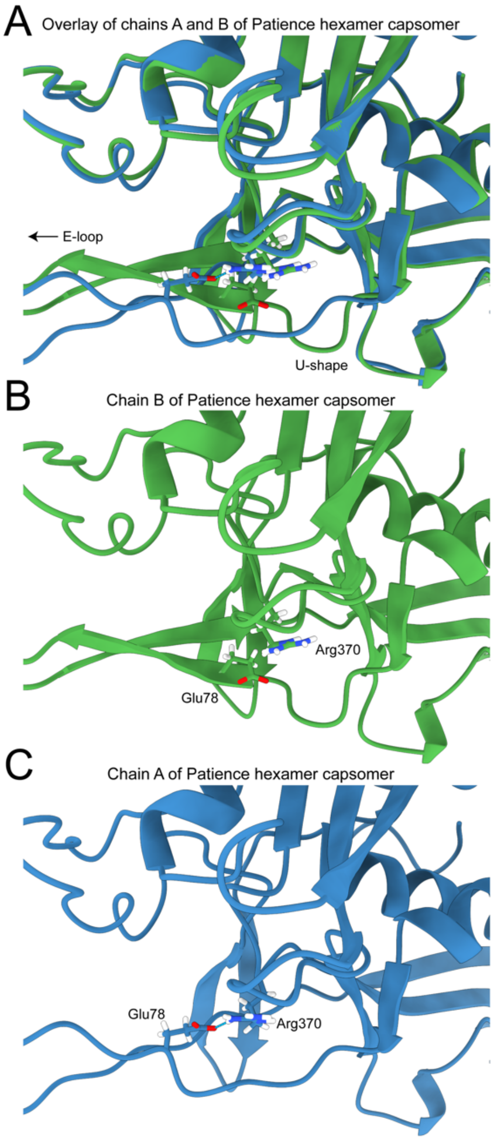


**Figure S6. Straightening of the E-loop of Chain A to bridge the 2-fold chasm.** Shows the overlay of chains A and B of the hexamer capsomer (A) as well as the two chains separately with Arg370 and Glu78 highlighted (B and C).

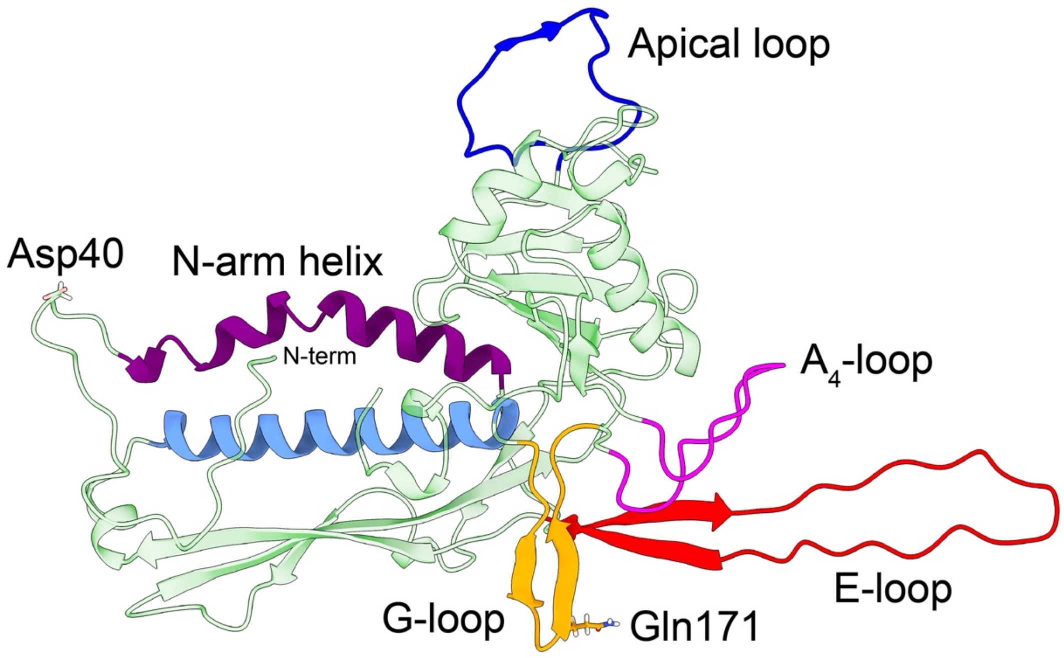


**Figure S7.** The hexamer major capsid protein.

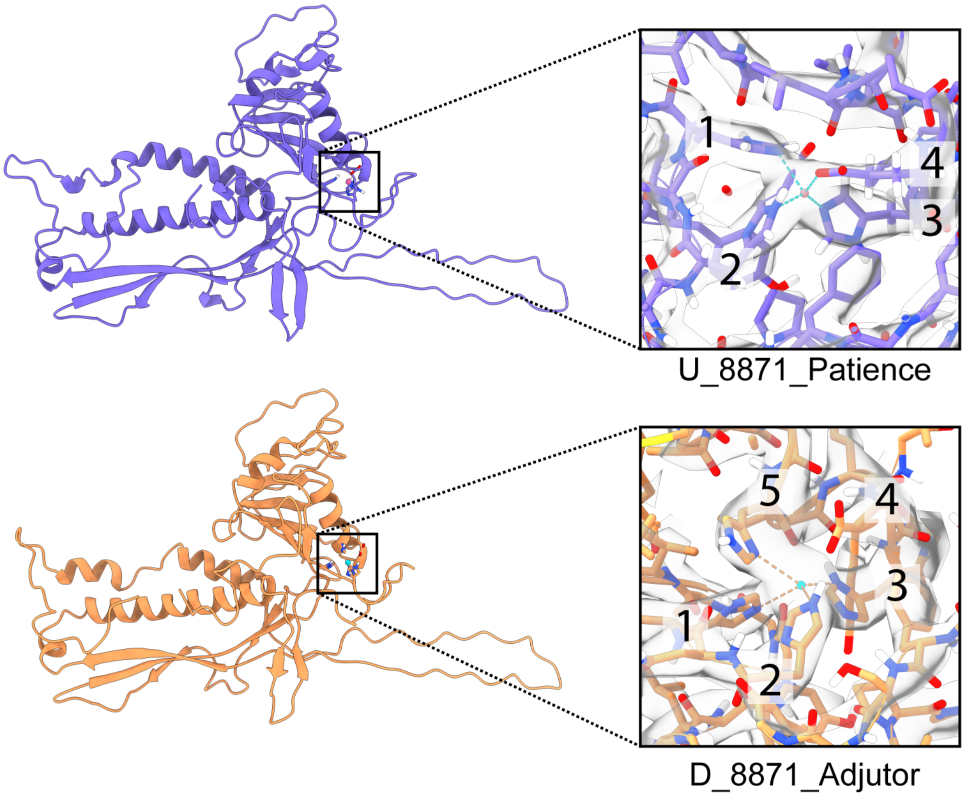


**Figure S8. Metal co-ordination in Patience and Adjutor.** The cryo-EM derived models are shown on the left with the putative metal binding domain highlighted with a black box. On the right the putative metal binding domain is shown closer up with the cryo-EM map overlaid (grey color). Predicted contacts are represented by dashed lines and residues potentially involved in the metal coordination are numbered.

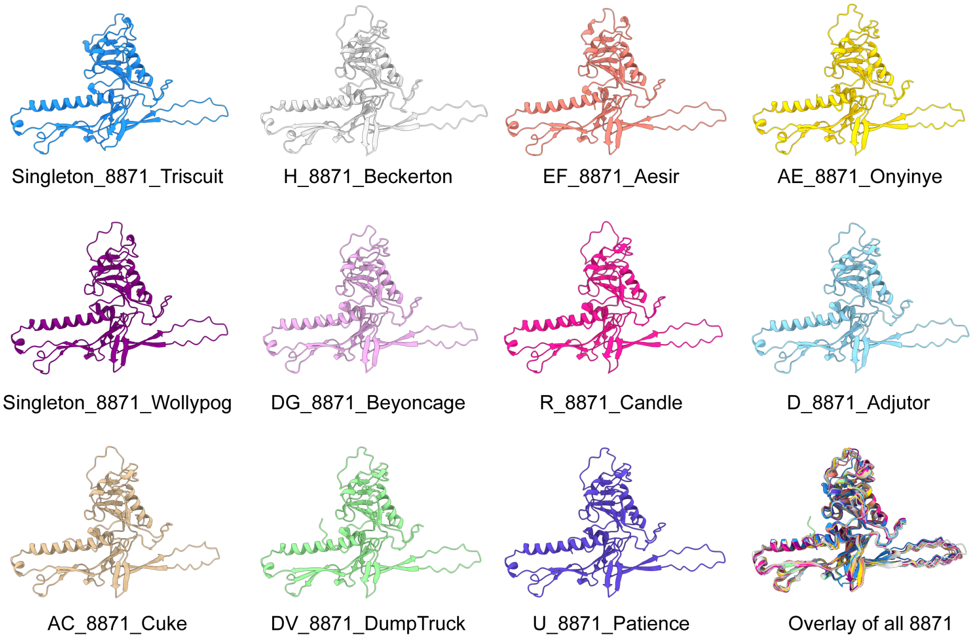


**Figure S9.** Predicted structures of all the major capsid proteins found in the Patience-like major capsid protein family. All structures were predicted with Alphafold. The N-terminus was truncated to where it crosses behind the spine helix due to the poor prediction.

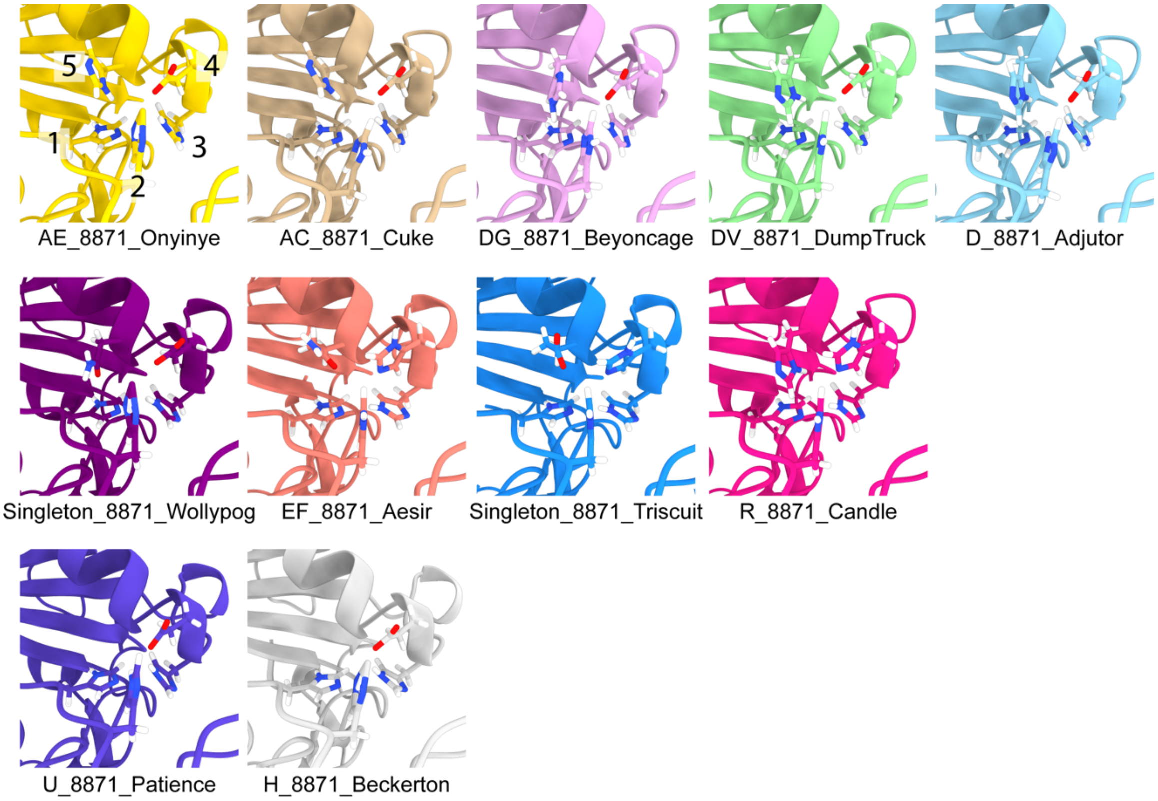


**Figure S10.** Metal co-ordination in the Patience-like major capsid protein. All structures shown were predicted with Alphafold. No metal ion has been modeleld in. Every major capsid protein of the Patience-like major capsid protein family shows similar metal co-ordination. In Onyinye, the five amino acids involved in the co-ordination are labelled. The first row all show similar co-ordination and have the same amino acids. The second row highlights the major capsid proteins that have different amino acids in positions 4 and 5. The final row shows Patience and Beckerton that lack the position 5 amino acid.

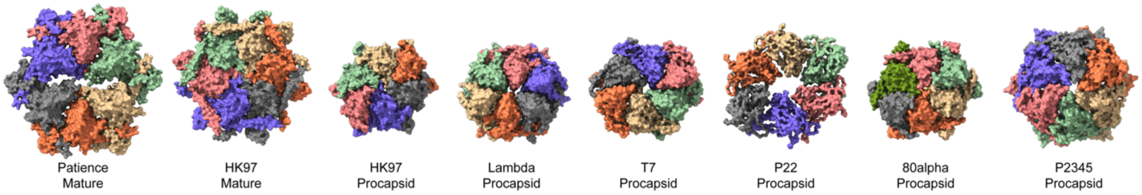


**Figure S11. HK97-fold procapsid hexamer structures.** Six procapsid hexamer structures with the mature capsid hexamer of Patience and HK97 bacteriophage for reference. Original PDB files are as follows. HK97 mature capsid (1OHG). HK97 procapsid (3E8K). Lambda procapsid (7VI9). P22 procapsid (2XYY). P2345 procapsid (6IBC). T7 procapsid (3J7V).

**Table S1.** Cryo-EM collection parameters, analysis, and final resolutions.

| Data collection | Adjutor | Patience |
|---|---|---|
| Microscope | Titan Krios | Titan Krios |
| Acceleration voltage (keV) | 300 | 300 |
| Spherical aberration (mm) | 2.7 | 2.7 |
| Pixel size / Å | 0.40075 | 0.3915 |
| Nominal defocus / μm | 0.8 to 2.1 | 1 to 3 |
| Detector (mode) | Gatan K3 (super resolution) | Falcon III (counting mode) |
| Total exposure dose / eÅ-2 | 30 | 30 |
| Number of frames | 30 | 30 |
| Number of micrographs | 6664 | 8640 |
| Number of particles in final refinement | 44239 | 61957 |
| Extract box size (fourier crop box size) | 2560/800 | 2400/800 |
| Final pixel size used in reconstruction | 1.28 | 1.19745 |
| Ewald sphere correction mask diameter | 760 | 788 |
| Symmetry | I (I1) | I (I1) |
| Resolution (FSC 0.143) | 2.66 | 2.39 |

**Table S2.** Members of the Patience-like major capsid protein family of Patience and Adjutor. Gp4 homologs are all putative and not confirmed by cryo-EM (apart from Patience and Adjutor that have been identified in this paper).

| Cluster | Number of members | Gp4 homolog pham | Average genome length (bp) | Host | Life Cycle |
|---|---|---|---|---|---|
| **AC** | 4 | 18383 | 70,029 | Mycobacterium | Lytic |
| **AE** | 2 | Unknown | 71,497 | Mycobacterium | Unknown |
| **D** | 21 | 6785 | 64,805 | Mycobacterium | Lytic |
| **DG** | 9 | Unknown | 66,155 | Gordonia | Lytic |
| **DV** | 17 | Unknown | 67,413 | Gordonia | Unknown |
| **EF** | 22 | Unknown | 56,496 | Microbacterium | Lytic |
| **H** | 10 | 16432 | 69,108 | Mycobacterium | Lytic |
| **R** | 8 | 36408 | 71,339 | Mycobacterium | Lytic |
| **U** | 3 | 16432 | 66,864 | Mycobacterium | Lytic |
| **Singleton_Triscuit** | 1 | Unknown | 67,539 | Microbacterium | Unknown |
| **Singleton_Wollypog** | 1 | Unknown | 63,364 | Arthrobacter | Unknown |

**Table S3.** Interactions between Adjutor gp4 and hexamer major capsid proteins. Interactions predicted with the PDBsum server.

| Chain | Number of interface residues | Interface area ($A^2$) | Number of salt bridges | Number of hydrogen bonds | Number of non-bonded contacts |
|---|---|---|---|---|---|
| **Gp4:A** | 27:33 | 1765:1598 | 4 | 17 | 183 |
| **Gp4:B** | 2:2 | 97:95 | 0 | 1 | 3 |
| **Gp4:C** | - | - | - | - | - |
| **Gp4:D** | - | - | - | - | - |
| **Gp4:E** | 4:5 | 257:233 | 0 | 1 | 9 |
| **Gp4:F** | 26:25 | 1598:1534 | 5 | 20 | 156 |

**Table S4.** Interactions between Patience gp4 and hexamer major capsid proteins. Interactions predicted with the PDBsum server.

| Chain | Number of interface residues | Interface area ($A^2$) | Number of salt bridges | Number of hydrogen bonds | Number of non-bonded contacts |
|---|---|---|---|---|---|
| **Gp4:A** | 47:55 | 2993:2755 | 1 | 27 | 267 |
| **Gp4:B** | 3:3 | 160:164 | 0 | 2 | 18 |
| **Gp4:C** | 1:1 | 66:59 | 0 | 1 | 6 |
| **Gp4:D** | 5:9 | 308:283 | 3 | 16 | 149 |
| **Gp4:E** | 14:22 | 818:704 | 0 | 5 | 72 |
| **Gp4:F** | 27:28 | 1481:1484 | 5 | 17 | 142 |

**Table S5.** Metal Ion-Binding site prediction scores of the putative metal binding sites in the Patience-like family of major capsid proteins. The Alphafold prediction pdb files were used with the online server.

| **Metal Ion** | **Patience (score/residues involved)** | **Adjutor (score/residues involved)** | **Candle (score/residues involved)** |
|---|---|---|---|
| **$Ca^{2+}$** | 0/None | 0/None | 0/None |
| **$Cu^{2+}$** | 0.933/185P, 198H, 200H, 224H | 0.963/202H,224H,228H | 1.035/207H,229H,233H |
| **$Fe^{3+}$** | 0.721/198H/200H/223E | 0.896/202H,204H,224H | 0.943/207H,229H,232H, 233H |
| **$Mg^{2+}$** | 0/None | 1.054/202H,204H | 1.150/207H,209H |
| **$Mn^{2+}$** | 0/None | 0/None | 0/None |
| **$Zn^{2+}$** | 1.077/198H,200H,226Y | 1.259/202H,204H | 1.214/207H,209H,229H |
| **$Cd^{2+}$** | 0/None | 0/None | 0/None |
| **$Fe^{2+}$** | 0.873/198H,200H,224H,372R | 1.156/202H,204H | 0.791/208D,209H,232H, 233H |
| **$Ni^{2+}$** | 1.059/198H,200H,223E | 0.966/202H,204H, 227E | 1.048/209H,229H |
| **$Hg^{2+}$** | 0.580/200H/377I | 0.739/198F,202H | 0.550/209H,387V |
| **$Co^{2+}$** | 1.276/198H,200H,223E | 1.098/224H,228H | 1.177/229H,233H |
| **$Cu^{2+}$** | 0.750/200H/220L/224H | 0.915/202H,204H,228H | 1.35/232H,233H |

**Table S6.** Geometrical analysis of the HK97-fold capsid range. The columns include the triangulation number (T), the hexagonal coordinates separating nearby pentamers (h,k), the class P, the multiplicity f, the generating hexagonal coordinates of the class, (h0,k0), the presence of a hexamer centered on the 3-fold axis (3-fold H), the triangulation number divided by three (T/3), and the hexagonal coordinates of the 3-fold hexamer ($h_H$, $k_H$). The capsids containing a hexamer centered on the 3-fold axis are shaded in grey.

| T | h | k | P | f | $h_0$ | $k_0$ | 3-fold H | T/3 | $h_H$ | $k_H$ |
|---|---|---|---|---|---|---|---|---|---|---|
| 1 | 1 | 0 | 1 | 1 | 1 | 0 | no | 1/3 | | |
| 3 | 1 | 1 | 3 | 1 | 1 | 1 | yes | 1 | 1 | 0 |
| 4 | 2 | 0 | 1 | 2 | 1 | 0 | no | 4/3 | | |
| 7 | 2 | 1 | 7 | 1 | 2 | 1 | no | 7/3 | | |
| 9 | 3 | 0 | 1 | 3 | 1 | 0 | yes | 3 | 1 | 1 |
| 12 | 2 | 2 | 3 | 2 | 1 | 1 | yes | 4 | 2 | 0 |
| 13 | 3 | 1 | 13 | 1 | 3 | 1 | no | 13/3 | | |
| 16 | 4 | 0 | 1 | 4 | 1 | 0 | no | 16/3 | | |
| 19 | 3 | 2 | 19 | 1 | 3 | 2 | no | 19/3 | | |
| 21 | 4 | 1 | 21 | 1 | 4 | 1 | yes | 7 | 1 | 2 |
| 25 | 5 | 0 | 1 | 5 | 1 | 0 | no | 25/3 | | |
| 27 | 3 | 3 | 3 | 3 | 1 | 1 | yes | 9 | 3 | 0 |
| 28 | 4 | 2 | 7 | 2 | 2 | 1 | no | 28/3 | | |
| 31 | 5 | 1 | 31 | 1 | 5 | 1 | no | 31/3 | | |
| 36 | 6 | 0 | 1 | 6 | 1 | 0 | yes | 12 | 2 | 2 |
| 37 | 4 | 3 | 37 | 1 | 4 | 3 | no | 37/3 | | |
| 39 | 5 | 2 | 39 | 1 | 5 | 2 | yes | 13 | 1 | 3 |
| 43 | 6 | 1 | 43 | 1 | 6 | 1 | no | 43/3 | | |
| 48 | 4 | 4 | 3 | 4 | 1 | 1 | yes | 16 | 4 | 0 |
| 49 | 5 | 3 | 49 | 1 | 5 | 3 | no | 49/3 | | |
| 49 | 7 | 0 | 1 | 7 | 1 | 0 | no | 49/3 | | |
| 52 | 6 | 2 | 13 | 2 | 3 | 1 | no | 52/3 | | |
